# A mouse model of hereditary hemorrhagic telangiectasia generated by transmammary-delivered immunoblocking of BMP9 and BMP10

**DOI:** 10.1101/084889

**Authors:** Santiago Ruiz, Haitian Zhao, Pallavi Chandakkar, Prodyot K. Chatterjee, Lionel Blanc, Christine N. Metz, Fabien Campagne, Philippe Marambaud

## Abstract

Hereditary hemorrhagic telangiectasia (HHT) is a potentially life-threatening genetic vascular disorder caused by loss-of-function mutations in the genes encoding activin receptor-like kinase 1 (ALK1), endoglin, Smad4, and bone morphogenetic protein 9 (BMP9). Injections of mouse neonates with BMP9/10 blocking antibodies lead to HHT-like vascular defects in the postnatal retinal angiogenesis model. Mothers and newborns share the same immunity through the transfer of maternal antibodies during breastfeeding. Here, we investigated whether the transmammary delivery route could improve the ease and consistency of administering anti-BMP9/10 antibodies in the postnatal retinal angiogenesis model. We found that anti-BMP9/10 antibodies, when intraperitoneally injected into lactating dams, are efficiently transferred into the circulation of breastfed neonatal pups. Strikingly, pups receiving anti-BMP9/10 antibodies *via* breastfeeding displayed consistent and robust vascular pathology in the retina, which included hypervascularization and defects in arteriovenous specification, as well as the presence of multiple and massive arteriovenous malformations. Furthermore, RNA-Seq analyses of neonatal retinas identified an increase in the key pro-angiogenic factor, angiopoietin-2, as the most significant change in gene expression triggered by the transmammary delivery of anti-BMP9/10 antibodies. Transmammary-delivered BMP9/10 immunoblocking in the mouse neonatal retina is therefore a practical, noninvasive, reliable, and robust model of HHT vascular pathology.

## INTRODUCTION

Hereditary hemorrhagic telangiectasia (HHT)—also known as Rendu-Osler-Weber syndrome—is an autosomal dominant genetic disease affecting approximately 1 in 5,000-10,000 individuals ^1,2^. HHT is characterized by the presence of vascular anomalies in multiple tissues and organs in the form of arteriovenous malformations (AVMs) and mucocutaneous telangiectases^3,4^. These vascular anomalies are structurally compromised and fragile, and hence are susceptible to rupture and hemorrhage. Telangiectases on the skin and in the nasal, oral, and gastrointestinal mucosa, as well as AVMs in the brain, lungs, and liver are typical in the clinical presentation of HHT ^3,4^. Genetic studies have revealed that most HHT patients carry mutations in the genes *ENG* (encoding endoglin) or *ACVRL1* (activin receptor-like kinase 1, ALK1), which define the two disease subtypes HHT1 and HHT2, respectively ^5^. Mutations in *SMAD4* (encoding Smad4) and *GDF2* (bone morphogenetic protein 9, BMP9) were also found to cause rare forms of juvenile polyposis/HHT combined syndrome and HHT-like vascular anomaly syndrome, respectively ^5^. Strikingly, BMP9, endoglin, ALK1, and Smad4 all functionally interact in, and are key mediators of, the same BMP receptor signaling pathway. ALK1 is a BMP type I Ser/Thr kinase receptor of the transforming growth factor-β superfamily, which forms functional complexes with a BMP type II receptor (e.g., BMPRII) and the co-receptor endoglin. BMP9 and BMP10 were identified as the specific and physiological ligands for ALK1 in endothelial cells ^6–9^. Upon binding to ALK1-endoglin receptors, BMP9 and BMP10 activate the phosphorylation of receptor-regulated Smads (Smad1, Smad5, and Smad9) to facilitate the formation of Smad1/5/9-Smad4 complexes, which translocate into the nucleus to act as transcription factors in specific gene expression programs ^10,11^.

The exact mechanisms driving HHT pathogenesis remain unknown. However, recent evidence strongly suggests that the disease is caused by abnormal activation of angiogenesis, a process causing excessive endothelial cell proliferation and hypervascularization, which ultimately leads to the development of AVMs ^12^. Indeed, ALK1 and endoglin are predominantly expressed by endothelial cells and BMP9/10 signaling is required for proper vascular development and maintenance, as well as for controlling transcriptional responses critically involved in angiogenesis, such as Notch and Wnt signaling pathways ^13–16^.

Studies in cell lines and animal models have demonstrated that HHT mutations cause haploinsufficiency. Indeed, transfection experiments showed that HHT mutations in either ALK1 or endoglin blocked activation of Smad1/5/9 signaling by BMP9 ^17–19^. The notion that HHT is caused by loss-of-function mutations was strengthened by studies in mouse models showing that *Acvrl1* or *Eng* deficiency, or injection of ALK1 extracellular domain-derived ligand trap (ALK1-Fc), led to vascular hyperproliferation and AVMs ^20^. It is important to note that the AVMs in these models were consistently and robustly observed when *Acvrl1* or *Eng* was completely deleted or inhibited, and when precipitating events were in place, such as angiogenesis or inflammation, a process referred to as the double or multiple hit hypothesis ^20,21^. Consistent with these observations, simultaneous neutralization of BMP9 and BMP10 in the postnatal retinal angiogenesis model [by injection of anti-BMP10 antibodies (Abs) into BMP9 knockout (KO) mice, or by co-injection of anti-BMP9 and anti-BMP10 Abs into wild type mice] led to hypervascularization ^22,23^, a vascular defect that was also observed in *Acvrl1* or *Eng* conditional KO models and thus is reminiscent of HHT pathology ^20^.

The postnatal retinal model is commonly used to study physiological or pathological angiogenesis during vascular development. Indeed, the newborn mouse retina is avascular and blood vessels rapidly develop during the first 2 weeks of life through angiogenesis ^24^. This developmental characteristic offers a unique opportunity to study and/or manipulate vascular development, maintenance, and remodeling. In order to improve the BMP9/10 neutralization retinal angiogenesis model, we tested whether the natural transmammary delivery route for administration of BMP9/10 blocking Abs could replace the stressful, cumbersome, and poorly reliable step of individually injecting large numbers of pups.

Neonatal immunity is controlled by passive immunoglobin transfer from the mothers to their newborns during breastfeeding. This mechanism of transmammary delivery of Abs can be very efficient because Abs survive the gastrointestinal tract of the neonates to be absorbed into the blood circulation ^25^. Here we report that BMP9 and BMP10 blocking Abs are efficiently transferred from the dam’s circulation into the blood of their breastfed neonates to generate a consistent and robust retinal vascular pathology, which was characterized by the presence of key HHT vascular defects. Thus, delivery of BMP9/10 blocking Abs to neonatal pups via breast milk produces a practical, noninvasive, reliable, and robust model of HHT vascular pathology.

## RESULTS

### Transmammary transfer of BMP9 and BMP10 blocking Abs into the circulation of breastfed mouse neonates

BMP9 and BMP10 are required, and have overlapping functions, during vascular development in the retina. Consequently, simultaneous injections of BMP9 and BMP10 blocking Abs directly into mouse neonates between postnatal day 1 (P1) and P4 was previously shown to induce strong retinal hypervascularization starting at P5 ^22,23^. In an effort to improve the ease and consistency of administering Abs to large cohorts of neonates, we investigated whether delivery *via* the transmammary route was suitable for BMP9/10 neutralization in the retinal angiogenesis model. In this context, we first asked whether BMP9 and BMP10 blocking Abs could transfer from the dams’ blood circulation to the blood of their breastfed neonates. To this end, lactating dams were injected intraperitoneally on P3 with monoclonal Abs specific for BMP9 (IgG2b, 15 mg/kg) and BMP10 (IgG2a, 15 mg/kg). As controls, separate sets of dams were injected with the same amount of isotype control IgG2b and IgG2a Abs, or with vehicle only (phosphate-buffered saline, PBS). At P6, neonates were euthanized and the levels of IgG2a and IgG2b Abs in their serum were analyzed by specific anti-IgG2a and anti-IgG2b ELISAs, respectively. We found that neonates breastfed by mice injected with either anti-BMP9/10 Abs or control IgGs showed a significant and consistent ~3-fold increase in serum IgG2a concentration (which corresponds to an elevation of ~40 μg/mL), when compared to PBS-treated controls (Fig. 1A). An elevation in IgG2b Abs in these neonate serum samples was not detected using the anti-IgG2b ELISA (Fig. 1B). Baseline serum concentrations for IgG2b Abs were of ~200 μg/mL (Fig. 1B), which was approximately 10-fold higher than baseline serum IgG2a levels (~20 μg/mL, Fig. 1A). We believe that these higher levels of baseline IgG2b in neonate serum could have masked any elevation of the concentration of this isotype after the transmammary transfer of IgGs. In this context, we developed an ELISA aimed at specifically detecting anti-BMP9 IgGs in the mouse serum (see Methods). Using this ELISA, we detected the presence of ~30 μg/mL of anti-BMP9 IgGs in the serum of neonates breastfed by dams injected with anti-BMP9/10 Abs, whereas, as expected, no anti-BMP9 IgG was measured in serum samples from control IgG-treated or PBS-treated neonates (Fig. 1C). Together, these results show that the transmammary route facilitated a robust and consistent transfer of both BMP9 and BMP10 blocking Abs to the neonatal circulation.

**Figure 1:**
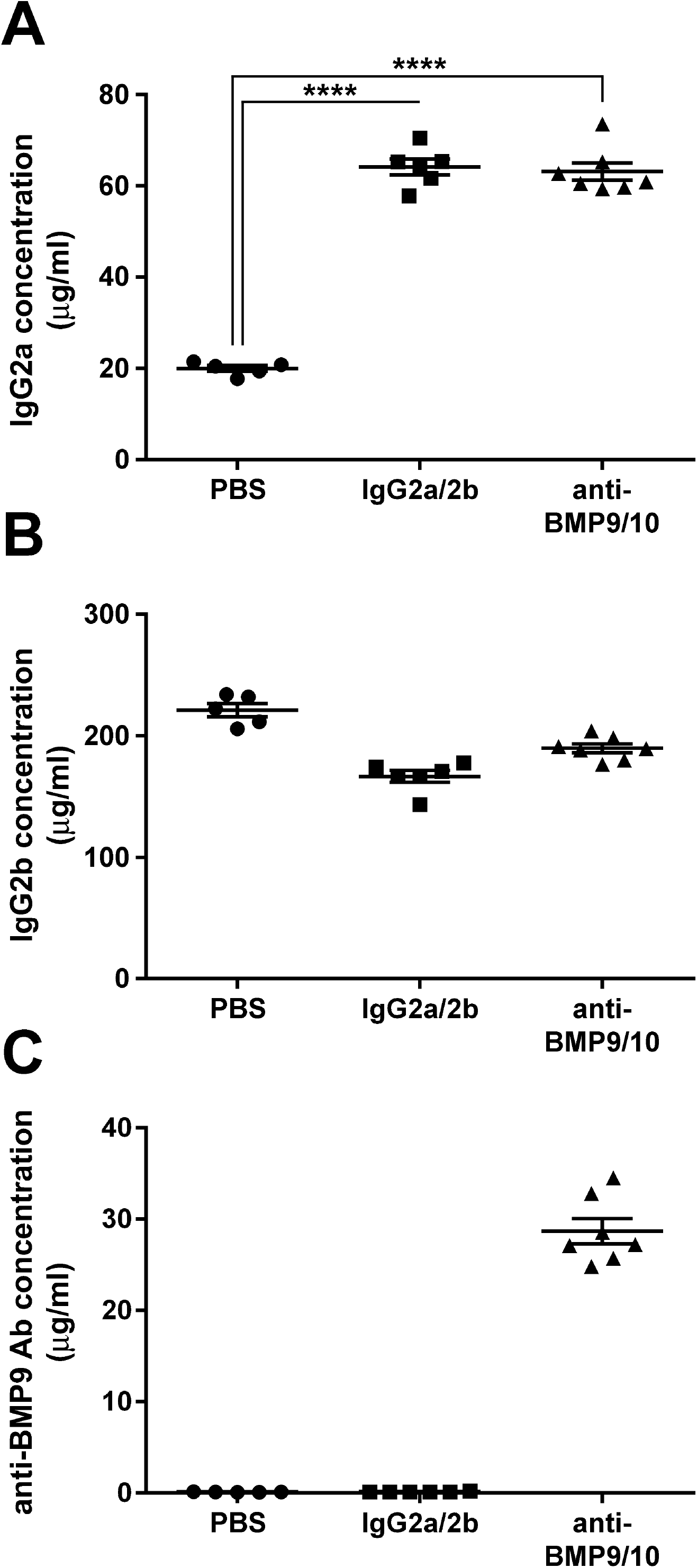
Transmammary transfer of BMP9 and BMP10 blocking Abs into the circulation of breastfed mouse neonates. (A-C) ELISAs were performed to measure IgG2a (A), IgG2b (B), and anti-BMP9 Ab (C) levels in the serum of P6 neonates breastfed by dams injected i.p. on P3 with either vehicle (PBS), isotype control IgGs (IgG2a/2b), or anti-BMP9/10 Abs. Data represents mean ± s.e.m. **** *P* < 0.0001, ANOVA.

### Transmammary transfer of BMP9 and BMP10 blocking Abs leads to abnormal hypervascularization in the neonatal retina

The retinal vasculature of P6 neonates was analyzed by histology techniques using isolectin B4 staining. Consistent with the data obtained after pup injections of anti-BMP9/10 Abs ^22,23^, a robust increase in postnatal retinal vascular density was observed after transmammary transfer of anti-BMP9/10 Abs, compared to retinas from control IgG-treated or PBS-treated neonates (Figs. 2A-C). Vessel staining revealed the clear presence of a hyperbranched vascular plexus characteristic of a defect in vessel patterning. We quantified the surface area occupied by the vasculature at the capillary plexus between arteries and veins (Figs. 2D-F), and at the arterial (Figs. 2G-I) and vascular (Figs. 2J-L) fronts. We found that, compared to retinas from control IgG-treated or PBS-treated neonates, anti-BMP9/10 Ab transfer significantly increased vascular density (Figs. 2M-O). In addition, the normal alternation of arteries and veins projecting from the optic disk (Figs. 2A and B) was compromised after anti-BMP9/10 Ab transfer (Fig. 2C). Consequently, some vessels could not be anatomically defined as veins or arteries (Fig. 2C), suggesting the presence of a defect in arteriovenous specification upon BMP9/10 immunoblocking.

**Figure 2:**
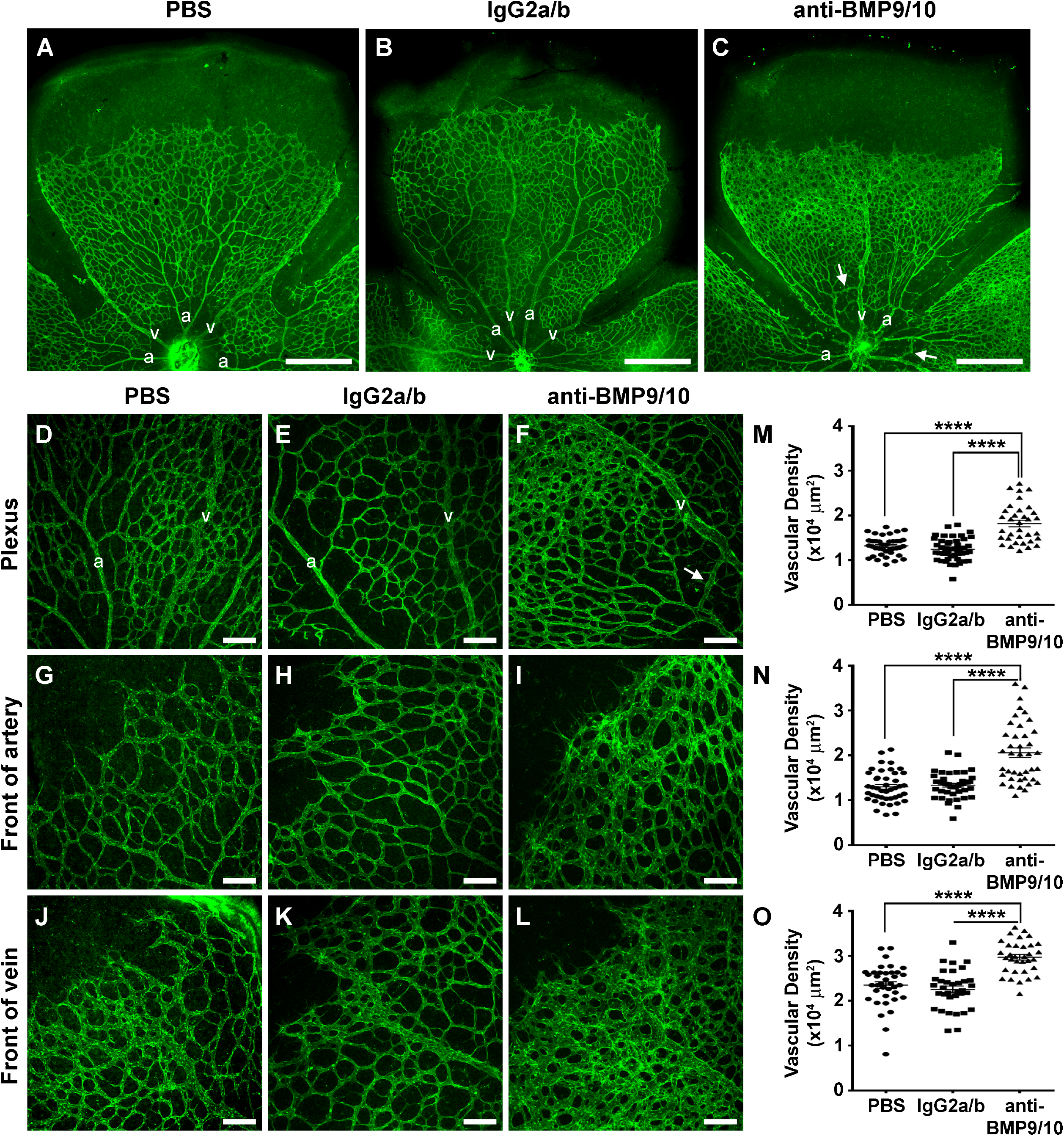
Transmammary transfer of BMP9 and BMP10 blocking Abs leads to abnormal hypervascularization in neonatal retinas. (A-C) Representative images of P6 fluorescent isolectin B4-stained retinas from neonate breastfed for 3 days by dams injected on P3 with PBS (A), control IgG2a/b Abs (B), or BMP9/10 blocking Abs (C). A, artery; V, vein; Scale bars, 500 μm. (D-L) Higher magnification showing retinal vasculature fields between an artery and a vein (Plexus, D-F), or at the front of an artery (G-I) or a vein (J-L). Scale bars, 100 μm. (M-O) Scatter plots showing the vascular density (n = 6 mice) between arteries and veins (M), or at the artery front (N) or vein front (O). Data represent mean ± s.e.m. **** *P* < 0.0001, ANOVA and Kruskal-Wallis test (M-O). Arrows in C and F indicate AVMs, defined as direct shunts between an artery and a vein.

### Transmammary transfer of BMP9 and BMP10 blocking Abs induces AVMs in the neonatal retina

Importantly, vessel staining using isolectin B4 also revealed the consistent presence of direct shunts between arteries and veins in neonatal retinas following anti-BMP9/10 Ab transmammary transfer, see arrows in Figs. 2C and 2F. To confirm the appearance of these AVMs, injections of latex dye in the circulation were used to better visualize pathological arteriovenous shunts in the retina. Because of its size, the latex dye cannot penetrate the capillary beds and thus, in a normal vasculature, is retained in the arterial circulation (see Ref. ^26^ and Fig. 3A for neonatal retinal artery visualization). However, under conditions of anti-BMP9/10 Ab transmammary transfer, the dye invaded the neonatal retinal vein circulation *via* multiple and, for some of them, very dilated AVMs (arrows, Fig. 3B). In order to quantify these arteriovenous defects, we measured the vascular arborization visualized by the latex dye. To this end, the Sholl’s concentric circle method of dendrite counting in neurons ^27^ was adapted to quantify the number of latex dye-positive vessels. Specifically, a grid of 5 concentric circles equidistantly drawn at 200 μm apart was superimposed at the center of the optic nerve (Fig. 3C) and vascular arborization was determined by counting the number of vascular crosses on each circle. A significant increase in the number of total retinal vascular crosses (Fig. 3D) and on the first 3 concentric circles (Fig. 3E) were found following anti-BMP9/10 Ab transmammary transfer, compared to control IgG-treated or PBS-treated retinas. Thus, anti-BMP9/10 immunoblocking using the transmammary route led to a significant vascular pathology in the retina, which included hypervascularization, defects in arteriovenous specification, and the presence of multiple and massive AVMs.

**Figure 3:**
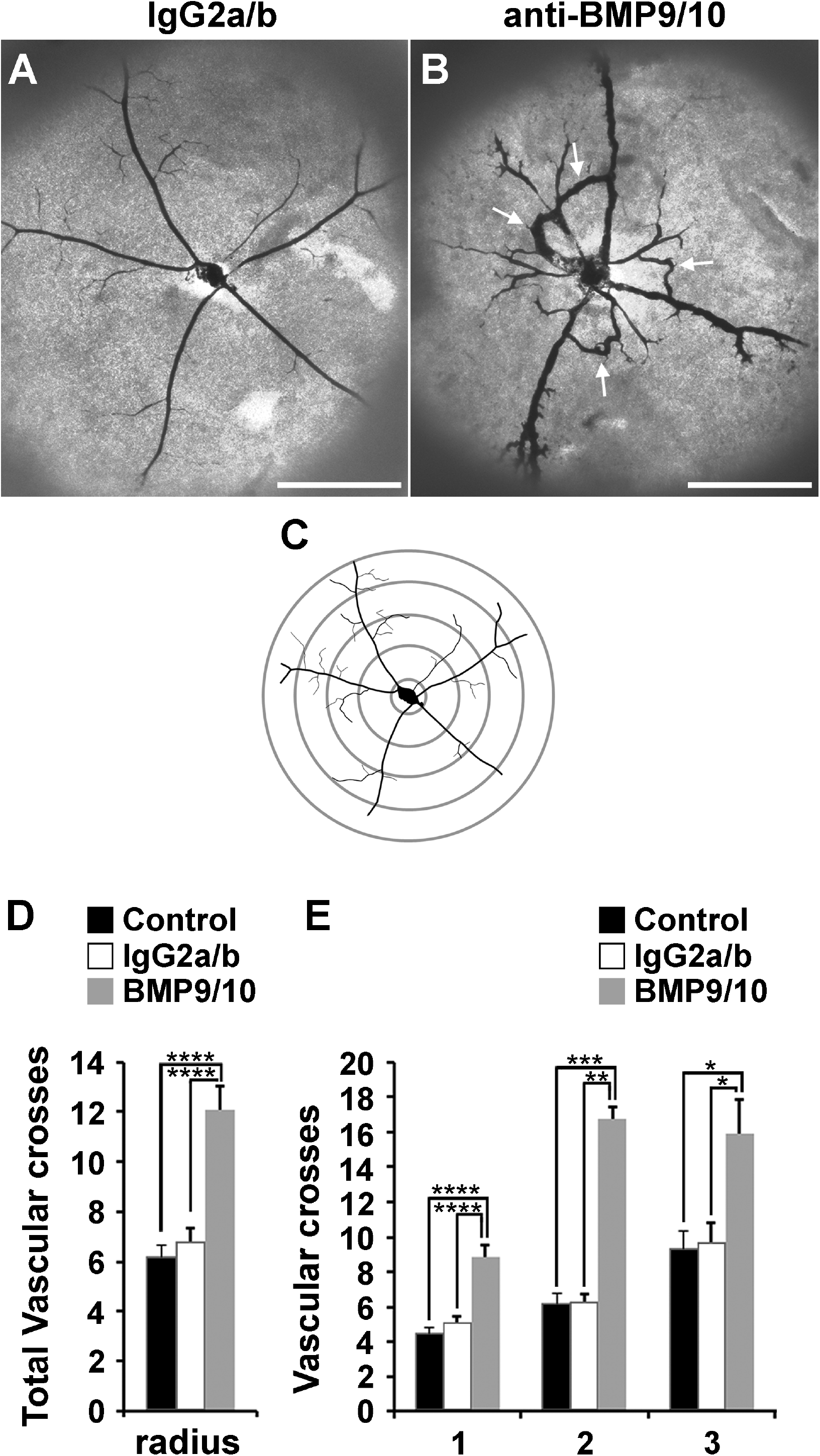
Transmammary transfer of BMP9 and BMP10 blocking Abs induces AVMs in neonatal retinas. (A-B) Representative images of blue latex-perfused retinal vasculature of P6 neonate breastfed for 3 days by dams injected on P3 with control IgG2a/b Abs (A) or BMP9/10 blocking Abs (B). Arrows in B indicate AVMs. Scale bars, 500 μm. (C) Scheme depicting the method employed for the quantification of the number of latex dye-positive vessels. (D) Histogram showing the total number of vascular crosses. (E) Histogram showing the number of vascular crosses per concentric circle (Control, n = 5; IgG2a/b, n = 7; anti-BMP9/10 Abs, n = 6 mice). Data represents mean ± s.e.m.; **** *P* < 0.0001, *** *P* < 0.001, ** *P* < 0.01, * *P* < 0.05; ANOVA and Kruskal-Wallis test (D and E).

### Gene expression changes in whole retinas and HUVECs after ALK1 signaling inhibition

Genome-wide transcriptomic analyses by RNA-Seq were employed to determine whether transmammary-delivered BMP9/10 immunoblocking affects gene expression in the whole retina. Gene expression profiles were analyzed in 6 retinas obtained from neonates following transmammary transfer of anti-BMP9/10 Abs or control IgG2a/2b Abs (n = 6 mice for each group). This screen identified 128 genes with significant expression changes (FDR ≤ 1%, absolute log2 fold change ≥ 0.7, Fig. 4A and Table S1). Notably, in BMP9/10-immunoblocked retinas, we found a significant increase in the expression of *Angpt2* (first hit when sorting by adjusted *P*-values, log2 fold change = 0.71, adjusted *P* = 2.69E-08, Limma Voom test, Table 1 and Fig. 4A). *Angpt2* codes for angiopoietin-2 (Ang2), a master regulator of sprouting angiogenesis ^28^ that was previously reported to be elevated at the transcriptional level in ALK1 deficient mice ^29,30^. Expression of genes coding for several collagen subunits, including *Col4a1* (log2 fold change = 0.70, adjusted *P* = 1.18E-06) and *Col15a1* (log2 fold change = 1.13, adjusted *P* = 5.05E-05), was also significantly increased in BMP9/10-immunoblocked retinas, compared to control retinas (Table 1 and Fig. 4A). Changes in collagen gene expression is of interest because collagen—and in particular type IV collagen—is a fundamental component of the vascular basement membrane and is key for endothelial cell proliferation and survival, and for angiogenesis ^31^. A function enrichment analysis, however, did not reveal any significant network that includes the identified deregulated genes [all categories with FDR > 0.5, GeneMania, ^32^].

**Table 1:**
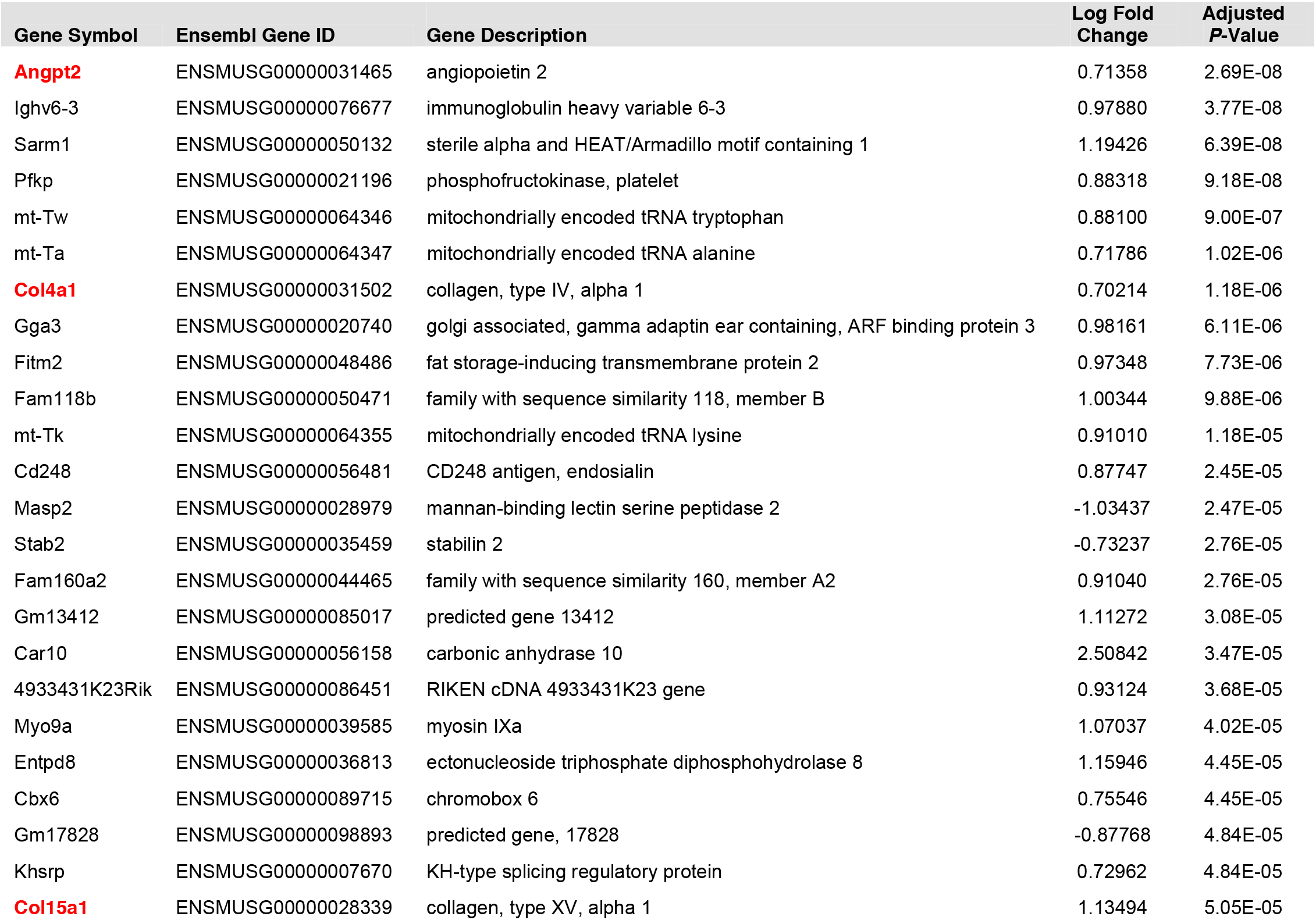
Top 25 retinal genes sorted by *P*-value, following transmammary-delivered immunoblocking of BMP9 and BMP10. The first 25 genes with adjusted *P*-value (false discovery rate) less or equal to 1%, and abs(logFC) ≥ 0.7, sorted by increasing adjusted *P*-value, are shown. Gene expression changes are expressed in log2 fold change and compare anti-BMP9/10 Ab-treated versus control IgG2a/2b-treated retinas. Positive logFC values denote genes over-expressed in anti-BMP9/10 Ab-treated retinas. A log2 fold change value of 1 corresponds to a doubling of expression in treated retinas (log2(2)=1).

**Figure 4:**
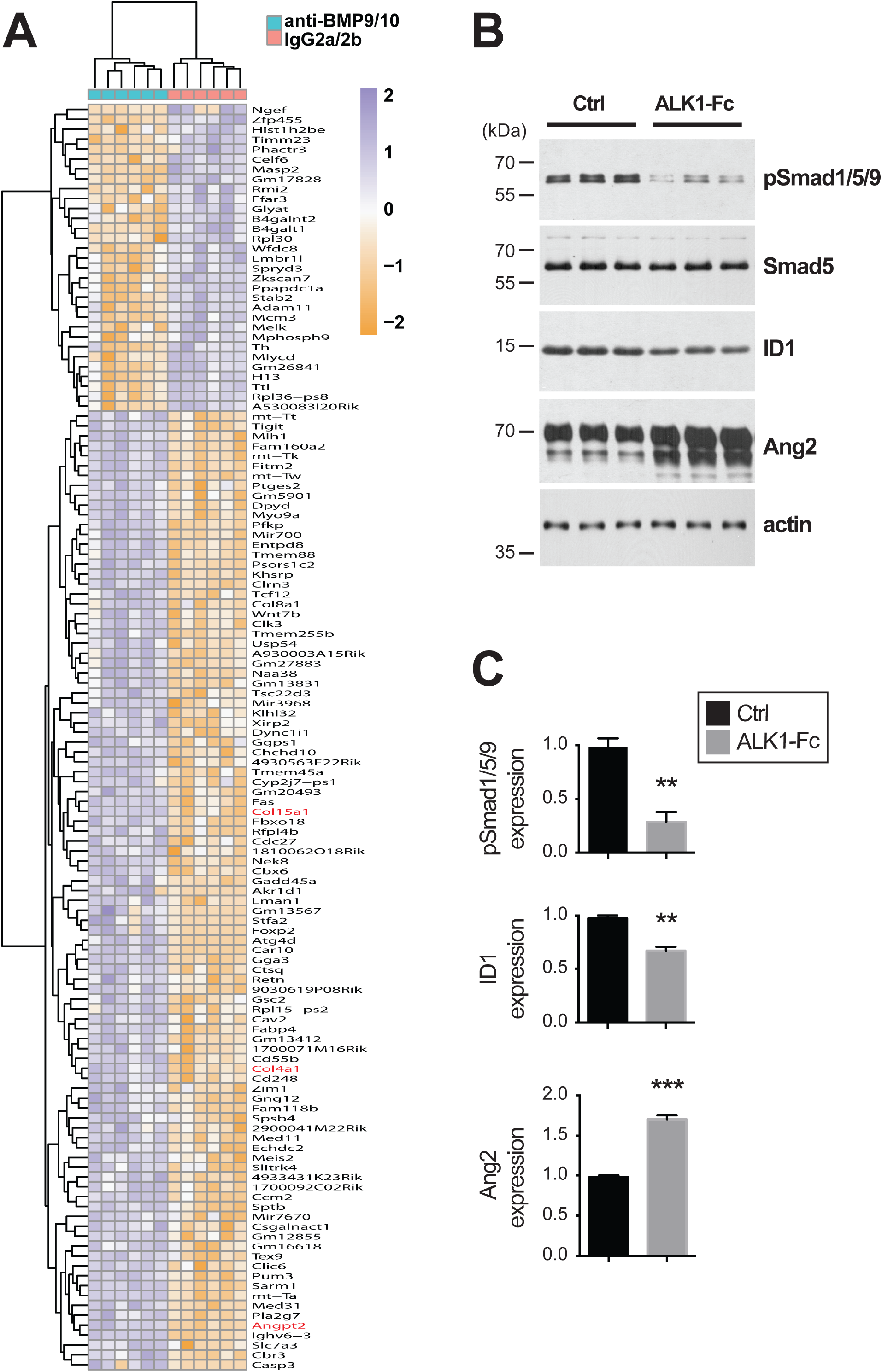
Gene expression changes in BMP9/10-immunoblocked retinas and ALK1-Fc-treated HUVECs. (A) Heat map displaying differently expressed genes in whole mouse retinas following transmammary transfer of anti-BMP9/10 or control IgG2a/2b Abs. (B) HUVECs were treated or not (Ctrl) with ALK1-Fc (1 μg/mL, 24 h). Cell extracts were then analyzed by WB using antibodies directed against the indicated proteins. (C) Densitometric analyses and quantification of phospho-Smad1/5/9, ID1, and Ang2 relative levels in three independent measurements as in (B). Results are mean ± s.e.m.; *** *P* < 0.001, ** *P* < 0.01 (unpaired Student’s t-test).

To confirm the effect of ALK1 inhibition on global gene expression and to determine which deregulations originate from endothelial cells, we also performed RNA-Seq on human umbilical vein endothelial cells (HUVECs) treated or not with the ligand trap ALK1-Fc. Comparison of the gene expression profiles obtained from BMP9/10-immunoblocked neonatal retinas and ALK1-Fc-treated HUVECs validated the increase of Ang2 expression upon ALK1 signaling inhibition. Indeed, ALK1-Fc treatment significantly increased of *ANGPT2* RNA expression in HUVECs (log2 fold change = 0.32, adjusted *P* = 2.99E-06). These results are important because they confirmed that ALK1 signaling is an endogenous repressor of Ang2 expression in endothelial cells. In order to confirm at the protein level the effect of ALK1 signaling inhibition on Ang2 expression, we analyzed Ang2 levels by western blot (WB) in ALK1-Fc-treated HUVECs. As expected, we found that ALK1-Fc treatment strongly suppressed ALK1 signaling, which was manifested by a decrease in the activating phosphorylation of Smad1/5/9 and in the protein levels of inhibitor of differentiation 1 (ID1), two downstream targets and effectors of ALK1 activation (Figs. 4B and C). Importantly and in line with the RNA-Seq data, we further found that ALK1-Fc treatment of HUVECs significantly increased Ang2 protein levels (Figs. 4B and C). Together these results show that, at the autonomous level in endothelial cells, ALK1 signaling negatively controlled the expression of the key pro-angiogenic mediator Ang2. Consequently, inhibition of ALK1 signaling in the presented model triggered a pathological pro-angiogenic response that led to hypervascularization and AVMs.

## DISCUSSION

The first HHT mouse models developed were constitutive *Acvrl1* and *Eng* KO mice. Based on the loss-of-function mutation paradigm in HHT, heterozygous KO mice for *Acvrl1* or *Eng* were expected to exemplify the most representative models for this disease. However, *Acvrl1*^+/-^ and *Eng*^+/-^ mice only displayed a mild phenotype with variable HHT-like vascular defects from one genetic background to another ^20^. More recently, powerful inducible conditional KO mice for *Acvrl1* or *Eng* were generated ^26,33,34^. Complete knockdown of either *Acvrl1* or *Eng* in these models led to more robust and reliable phenotypes, which included the appearance of HHT-like AVMs ^20^. It is important to note that in these models, complete *Acvrl1*- or *Eng*-deficiency *per se* was found not to be sufficient for generating the observed vascular defects and that additional triggers in the form of inflammatory and/or angiogenic stimuli were required at the sites where AVMs developed ^20,21,26^. Together with the observation that homozygous *Acvrl1* or *Eng* KO mice die at an early embryonic stage most likely from angiogenesis defects, these data highlight the existence of a clear interplay between angiogenesis and ALK1-endoglin signaling. Evidence is indeed strong to indicate that ALK1-endoglin signaling is necessary for the proper expression of angiogenesis in certain tissues, and that angiogenic stimuli, in return, can provide a trigger for the development of HHT phenotypes in models defective for ALK1-endoglin signaling. For instance, treatments with the master initiator of sprouting angiogenesis VEGF ^35^ facilitated the development of vascular dysplasia in *Acvrl1* and *Eng* KO mice ^36,37^, and VEGF neutralization using bevacizumab was found to be very effective at interfering with AVM development and progression in the wound-induced AVM mouse model ^38^. These observations are in line with several early-phase clinical trials supporting that bevacizumab treatments—using topical, submucosal, or i.v. infusion routes—were effective at reducing epistaxis in HHT patients ^39–41^.

In this context, the postnatal retinal angiogenesis model is very attractive for studying HHT vascular pathology. Indeed, physiological angiogenesis, which is required for postnatal vascular development, will provide the required additional trigger for vascular pathology development in mice deficient for ALK1 or endoglin expression ^33,42^ or in mice treated with BMP9/10 blocking Abs ^22,23^. An important advantage of this model is the rapid pathology occurrence during the first week of postnatal vascular development. In addition, this model limits handling of newborn mice and their invasive manipulation. The use of BMP9/10 blocking Abs provides an additional advantage—over *Acvrl1* or *Eng* KO mice—which is that Abs can be injected into any mouse strain, avoiding the phenotypic variability sometimes observed between mutant mice of different and complex genetic backgrounds ^20^.

For these reasons and in the current study, we sought to improve the practicality and reliability of the BMP9/10 immunoblocking neonatal retinal angiogenesis model. Based on strong supporting literature showing that neonatal immunity is controlled by passive immunoglobin transfer from mothers to their newborns during breastfeeding, we asked whether the transmammary route could be used to deliver BMP9/10 blocking Abs into neonatal mice. We found for the first time that these Abs were very efficiently transferred from the dam’s circulation to the neonates’ blood during breastfeeding. In addition, this transfer triggered the development of a robust and consistent retinal vascular pathology, which included hypervascularization, partial loss of arteriovenous specification, and most importantly, AVMs.

Using whole transcriptome analyses in the neonatal retina, we further revealed in this study a significant elevation of Ang2 expression following anti-BMP9/10 Ab transmammary delivery. Whole transcriptome analyses of ALK1-Fc-treated HUVECs confirmed the effect of ALK1 signaling inhibition on Ang2 expression and also showed that ALK1 inhibition increased Ang2 expression in an endothelial cell-autonomous manner. WB analyses of ALK1-Fc-treated HUVECs confirmed that inhibiting ALK1 signaling significantly increased Ang2 protein levels. Together, these data demonstrate that ALK1 signaling is a physiological repressor of Ang2 expression in endothelial cells. This observation is consistent with previous data showing that Ang2 expression is increased in ALK1^-/-^ embryos ^29^ and adult ALK1^+/-^ lung tissue ^30^, as well as in conditional ALK1 KO mouse models of central nervous system AVMs ^43^. Increases in Ang2 expression were also reported in the plasma and plexiform lesions of patients affected by pulmonary arterial hypertension (PAH), a condition that can be caused by mutations in ALK1 or BMPRII ^44^. These data, however, contrast with the observation that Ang2 levels might be reduced in the plasma of a small cohort of HHT2 patients, but not in HHT1 patients ^45^. In this context, our data strengthen the concept that Ang2 might be a crucial trigger for the vascular pathology caused by ALK1 signaling loss-of-function, and beyond that, for HHT pathogenesis. Ang2 is a circulating ligand that, in conjunction with VEGF signaling, has critical pro-angiogenic properties during the induction of vascular sprouting ^46^. Knowing that Ang2 is currently being investigated as a potential target for anti-angiogenic tumor therapy ^47^, it is tempting to speculate that anti-Ang2 approaches might also have therapeutic value in HHT.

A gene that also showed deregulated expression, albeit with less consistency inside the groups, was *Ccm2* (log2 fold change = 2.44, adjusted *P* = 1.72E-04, see Table S1). *Ccm2* is of interest because it codes for malcavernin, a protein that mediates interactions between blood vessel cells. Mutations have been identified in the orthologous *CCM2* human gene in patients with familial cerebral cavernous malformation (CCM) ^48^. CCM leads to vascular malformations mostly in the brain, but also in other tissues, and its phenotype is closely related to, albeit different from, the AVMs observed in HHT patients: CCMs are low-flow lesions, while HHT AVMs are high-flow lesions ^49^. Furthermore, while loss-of-function mutations in *CCM2* are associated with CCM in humans, in our model of BMP9/10-immunoblocking, we observed an increase of *Ccm2* expression, suggesting the possible existence of a compensatory mechanism that mitigates the vascular changes induced by ALK1 signaling inactivation. Further studies will be required to address this possible mechanism and determine whether *CCM2* deregulations might also be associated with AVM development in HHT.

In conclusion, we propose that transmammary-delivered immunoblocking of BMP9 and BMP10 in the mouse neonatal retina is a practical, noninvasive, reliable, and robust model to study HHT pathogenesis. Furthermore, this study provides proof-of-concept data that the described model is suitable for mechanistic and ultimately, pre-clinical therapeutic investigations for HHT. From a more general standpoint and to the best of our knowledge, the paradigm described here represents the first example of generation of an infection-unrelated disease model utilizing mother/neonate-shared immunity.

## MATERIALS AND METHODS

### Mice

All animal procedures were performed in accordance with protocols approved by the Feinstein Institute for Medical Research Institutional Animal Care and Use Committee. Timed-pregnant C57BL/6J mice (3-4 month old) were used in this study (The Jackson Laboratory).

### Antibody (Ab) injections and transmammary transfer of antibodies *via* lactation

Lactating dams were injected i.p. on P3 with PBS, mouse monoclonal isotype control Abs (15 mg/kg, IgG2b, MAB004; 15 mg/kg, IgG2a, MAB003; R&D Systems), or mouse monoclonal anti-BMP9 and anti-BMP10 Abs (15 mg/kg, IgG2b, MAB3209; 15 mg/kg, IgG2a, MAB2926; R&D Systems, respectively). Breastfed neonates were euthanized by CO_2_ asphyxiation on P6 and non-heparinized blood was collected. Neonates were enucleated and eyes fixed in 4% paraformaldehyde for 20 min on ice and retinas were isolated.

### Measurements of antibody levels by ELISA in neonatal mouse serum

ELISAs were used to measure IgG2a, IgG2b, and anti-BMP9 Ab levels in mouse serum. IgG2a and IgG2b ELISAs were performed as per the manufacturer’s instructions (eBioscience). For the anti-BMP9 Ab ELISA, the wells of 96-well ELISA plates (Maxisorp, Nunc) were coated with 100 μL of 1 μg/mL recombinant BMP9 (R&D Systems) in coating buffer (15 mM K_2_HPO_4_, 25 mM KH_2_PO_4_, 0.1 M NaCl_2_, 0.1 mM EDTA, 7.5 mM NaN_3_) and incubated overnight at 4^o^C. Plates were then washed 3 times with 0.05% Tween PBS (PBST) and blocked for 1 h at room temperature (RT) with 1% BSA in PBS. After washing 3 times with PBST, serial dilutions of individual mouse serum samples and reference mouse anti-BMP9 antibody (MAB3209, R&D Systems) (diluted in 1% BSA PBS) were prepared and 100 μL/well were incubated for 2 h at RT. After 3 more washes, 100 μL/well horseradish peroxidase (HRP)-conjugated goat anti-mouse immunoglobins (Igs) secondary antibody (Southern Biotech, diluted 1:500 in 1% BSA PBS) was incubated for 1 h at RT. TMB substrate was added after 5 washes and the reaction was allowed to develop for 30 min at RT. The optical density was measured at 450 nm using a TECAN GENios Pro plate reader.

### Retinal whole mount immunohistochemistry (IHC)

IHC was performed using the previously described protocol ^50^ with the following modifications. After fixation using 4% paraformaldehyde, retinas were dissected, cut four times to flatten them into petal flower shapes, and fixed with methanol for 20 min on ice. After removing methanol, retinas were washed in PBS for 5 min on a shaker at RT, and blocked in blocking solution (0.3% Triton, 0.2% BSA in PBS) for 1 h on a shaker at RT. Retinas were then incubated in isolectin GS-IB4 Alexa Fluor 488 (I21411, Molecular Probes) diluted 1:100 in blocking solution on a shaker overnight at 4°C. Retinas were then washed four times in 0.3% Triton in PBS for 10 min on a shaker, followed by two washes in PBS for 5 min on a shaker before mounted with Vecta Shield (H-1000, Vector Laboratories).

### Blue latex perfusion assay

The blue latex perfusion assay was performed using a previously described protocol ^26^, with the following modifications. Briefly, after euthanasia, neonate thoraxes were opened using Dumont #5 forceps and extra fine Bonn scissor (Fine Science Tools Inc.), the right atriums were cut using a Vannas spring scissor (Fine Science Tools Inc.), and the left ventricles were injected manually with 1 mL Blue latex (BR 80B, Connecticut Valley Biological Supply) using an insulin syringe U-100 (329652, BD Biosciences). After perfusion, eyes were enucleated and fixed, and retinas were dissected for microscopic observation as described above.

### Image acquisition and analysis

Images for the analysis of the vascular network density were acquired using a laser confocal microscopy Olympus FV300, while images for the analysis of the number of crosses of arteries and veins were acquired using a Nikon Eclipse T2000-S microscope. Quantifications were performed using ImageJ. The analysis of the vascular network density was performed on a total of 6 mice per condition (PBS, isotype control, and anti-BMP9/10 Abs). Using a 20x lens, images (2-5 fields per retina) were acquired in three different locations of the vasculature: The plexus (between an artery and a vein) and the front of the extending arterial and venous vasculature. Quantification was done using the measure particles tool, working with 8-bit images, adjusting the threshold, and measuring the endothelial area occupy by the vasculature in a region of interest of 200 x 200 μm^2^ in these three locations. For the analysis of the number of artery and vein crosses, whole retinas were imaged with a 4x lens (isotype control condition, n = 7 mice; anti-BMP9/10 Ab condition, n = 6 mice). Quantification was performed using the counter tool and plug-in concentric circles, counting each crossing artery and vein on 5 concentric circles equidistantly drawn (200 μm apart) from the optic nerve to the periphery of the extending vasculature of the retina.

### Statistical analysis

ANOVA was used to assess statistical significance within multiple comparisons analyses. When data had non-normal distribution, Kruskal-Wallis test was performed instead. Tukey’s or Dunn’s multiple comparisons test were performed as post hoc tests. In all cases GraphPad Prism version 6.0 was used.

### Retinal RNA extraction and sequencing

After euthanasia, neonate eyes were enucleated and retinas dissected in PBS. Retinas were rapidly dried with absorbent paper, snap-frozen in liquid nitrogen and stored at −80°C. RNA was then extracted from retinas using the RNeasy Micro Kit (Qiagen) according to the manufacturer’s instructions. Total RNA quality was verified using Thermo Scientific NanoDrop and Agilent Bioanalyzer. RNA was then processed for RNA-Seq at the Genomics Resources Core Facility, Weill Cornell Medical College, New York, NY. Briefly, cDNA conversion and library preparation were performed using the TrueSeq v2 Illumina library preparation kit, following manufacturers’ recommended protocol. Samples were multiplexed 6 per lane and sequenced on an Illumina HiSeq 4000 instrument.

### RNA-Seq data analysis

Reads were uploaded to the GobyWeb system ^51^ and aligned to the 1000 genome human reference sequence ^52^ with the STAR aligner ^53^. Ensembl annotations for transcripts were obtained from Biomart and Ensembl automatically using GobyWeb. Annotations were used to determine read counts using the Goby alignment-to-annotation-counts mode ^54^, integrated in GobyWeb in the differential expression analysis with EdgeR plugin. Counts were downloaded from GobyWeb as a tab delimited file and analyzed with MetaR ^55^. Statistical analyses were conducted with Limma Voom ^56^, as integrated in MetaR 1.7.2, using the rocker-metar docker image version 1.6.0.1. *P*-values were adjusted for multiple testing using the False Discovery Rate method ^57^. Heat maps were constructed with MetaR, using the pheatmap R package. Gene annotations were determined with Ensembl/Biomart, using the biomart micro-language in MetaR ^55^. MetaR analysis is shown in supplementary material.

### HUVEC cultures, treatments, RNA extraction, and western blot (WB) analyses

HUVECs were isolated from anonymous umbilical veins, as described before ^58^. HUVECs were subcultured using a trypsin/EDTA-reagent pack (Lonza) and maintained in endothelial cell growth media (Sciencell). Cells were treated or not for 24 h with 1 μg/mL ALK1-Fc (R&D Systems). Cells were then rinsed with PBS and processed for RNA extraction and RNA-Seq as described above. For WB analyses, cells were processed as before ^59^ with the following modifications. Cells were solubilized in RIPA buffer (EMD Millipore) supplemented with 1× Complete protease inhibitor mixture (Roche Applied Science). 5-20 μg of proteins (depending on the primary antibody used) were separated by SDS-PAGE and transferred onto nitrocellulose membranes. Membranes were then probed with Abs directed against phospho-Smad1/5/9 (Cell Signaling Technology), total Smad5 (Cell Signaling Technology), ID1 (BioCheck), Ang2 (Santa Cruz Biotechnology), and actin (BD Transduction Laboratories). A standard ECL detection procedure was then used.

## AUTHOR CONTRIBUTIONS

S.R., H.Z., P.C., and P.K.C. performed the experiments and analyzed the data. F.C. performed the bioinformatic analyses of the RNA-Seq data. P.M. conceived the project and elaborated the experimental strategy with S.R., L.B., C.N.M., and F.C. P.M., S.R., F.C., L.B., C.N.M. wrote the manuscript.

## ACCESSION CODES

RNA-Seq reads have been deposited to the Sequence Read Archive under accession code SRP071883.

